# spicyR: Spatial analysis of *in situ* cytometry data in R

**DOI:** 10.1101/2021.06.07.447307

**Authors:** Nicolas P. Canete, Sourish S. Iyengar, James S. Wilmott, John T. Ormerod, Andrew N. Harman, Ellis Patrick

## Abstract

**Motivation:** High parameter histological techniques have allowed for the identification of a variety of distinct cell types within an image, providing a comprehensive overview of the tissue environment. This allows the complex cellular architecture and environment of diseased tissue to be explored. While spatial analysis techniques have revealed how cell-cell interactions are important within the disease pathology, there remains a gap in exploring changes in these interactions within the disease process. Specifically, there are currently no established methods for performing inference on cell localisation changes across images, hindering an understanding of how cellular environments change with a disease pathology.

**Results:** We have developed the spicyR R package to perform inference on changes in the spatial localisation of cell types across groups of images. Application to simulated data demonstrates a high sensitivity and specificity. We demonstrate the utility of spicyR by applying it to a type 1 diabetes imaging mass cytometry dataset, revealing changes in cellular associations that were relevant to the disease progression. Ultimately, spicyR allows changes in cellular environments to be explored under different pathologies or disease states.

**Availability and Implementation:** R package freely available at http://bioconductor.org/packages/release/bioc/html/spicyR.html and shiny app implementation at http://shiny.maths.usyd.edu.au/spicyR/

**Supplementary information:** Code for reproducing key figures available at https://github.com/nickcee/spicyRPaper.

## Introduction

Identifying changes in the spatial distribution of cells is vital for understanding the cellular processes that are present in diseased tissue. Multiplexed histological techniques have allowed the complex cellular architecture and environment of diseased tissue to be explored by enabling the simultaneous profiling of multiple cell types. Fluorescence-based methods, including co-detection by imaging (CODEX; Goltsev et al. 2018), cyclic immunofluoresecence (cycIF; Lin et al. 2015), and iterative indirect immunofluorescence imaging (4i; Gut et al. 2018), as well as mass cytometry imaging techniques, including imaging mass cytometry (IMC; Giesen et al. 2014) and multiplexed ion beam imaging by time of flight (MIBI-TOF; Angelo et al. 2014) allow up to 40 protein markers to be visualised with single cell resolution. Additionally, the development of spatial-based transcriptomic techniques such as High-Definition Spatial Transcriptomics (Vickovic et al. 2019) and sequential fluorescence in situ hybridisation (seqFISH; Lubeck et al. 2014) allow tens of thousands of transcripts to be spatially resolved within an image at single-cell resolution. This large increase in the scale and dimensionality of the images being acquired has necessitated the development of analysis techniques capable of interrogating such complex data.

Established image analysis techniques have enabled the investigation of cell-cell interactions and cell migration within an image, facilitating the interrogation of high parameter imaging data in a single-cell manner. Standard pipelines start by identifying cells through single-cell segmentation followed by cell type classification by clustering or manually gating marker expression (Baharlou et al. 2019; Carpenter et al. 2006; Sommer et al. 2011; Van Valen et al. 2016). From here, a differential analysis of cell composition or marker expression within the image dataset can be performed to identify high-level associations with a phenotype of interest. The spatial dimension afforded by imaging can furthermore allow the spatial context of these cells to be quantified in multiple ways. One way to interrogate the spatial organization of cells is to quantify the spatial attraction or avoidance between pairs of cell types. This often involves counting an association measure between cell types. Counting the number of touching cells of a pair of cell types provides a measure of spatial association (Schapiro et al. 2017). Randomising the labels of cells can then be used to identify how significant a pairwise interaction is within an image, as seen in an application to type-1 diabetes images in Damond et al. (2019). This approach can be extended by calculating distances between cells and tabulating their nearest neighbours as used by Keren et al. (2018) to assess the spatial interactions involved in triple-negative breast cancer pathology. The use of summary functions such as Ripley’s K function (Ripley 1976, Baddeley et al. 2015), can further be used to assess how localisation between cell types vary with distance by modelling cells as point process. Further to this, identification of spatial communities (Jackson et al. 2020) have been performed through graph-based techniques, associating the spatial distribution of groups of cell types to disease outcomes. Finally, techniques such as spatial variance component analysis (Arnol et al. 2019) have been developed to identify the sources of variation of gene or protein markers attributed to cell-cell interactions. Overall, such techniques can identify spatial structure within high parameter images. This could then be attributed to the pathology of the disease states being studied.

A key gap that remains in the analysis of high parameter images is the identification of differential cell type localisation across groups – that is, changes in the extent of pairwise cell type localisation across these groups. These groups could be a clinical outcome such as disease stage or response to treatment or come from a perturbed experiment. Present strategies involve comparing association measures across groups. Damond et al. (2019) compares the number of each pairwise interaction using a Mann-Whitney’s U test to show changes in cell localisations across different disease stages. Farkkila et al. (2020) compares the Z-scores of the pairwise cell interaction obtained from bootstrapping across ovarian cancer clinical outcomes to identify changes in cell distribution. While such approaches are appropriate, they do not allow for the modelling of multiple images from multiple subjects. Additionally, these strategies ignore the variation observed in the quantification of cell type localisation.

In this manuscript we present an R package “SPatial analysis of In situ CYtometry data in R” (spicyR) and a corresponding web application to facilitate inference on changes in spatial localisations between cell types. spicyR aims to provide an easy-to-use approach to identify differential cell localisation analysis with respect to a disease or treatment and has the capacity to model information from multiple images per subject or account for images having a substantial difference in cell number. We demonstrate the performance of spicyR through simulation and apply the package to the diabetes imaging data presented in Damond et al. (2019), revealing changes in cellular localisation with type-1 diabetes progression.

## Materials and Methods

Our R package, spicyR, provides the framework for performing inference on the changes in spatial localisations between pairs of cell types which can be associated with a discrete or continuous outcome (**Fig. 1A**). As described below, spicyR consists of three primary steps: 1) summarising the degree of spatial localisation between pairs of cell types for each image; 2) modelling the variability in the localisation summary statistics as a function of cell counts, and 3) testing for changes in localisation associated with a response variable. The significance of this change is assessed using a linear model, or a mixed-effects model if there are multiple images belonging to a subject (**Fig. 1B**). The R package is available on Bioconductor, http://bioconductor.org/packages/release/bioc/html/spicyR.html, and is also implemented as an interactive shiny application http://shiny.maths.usyd.edu.au/spicyR/ for individuals who don’t want to use R.

**Figure 1.**
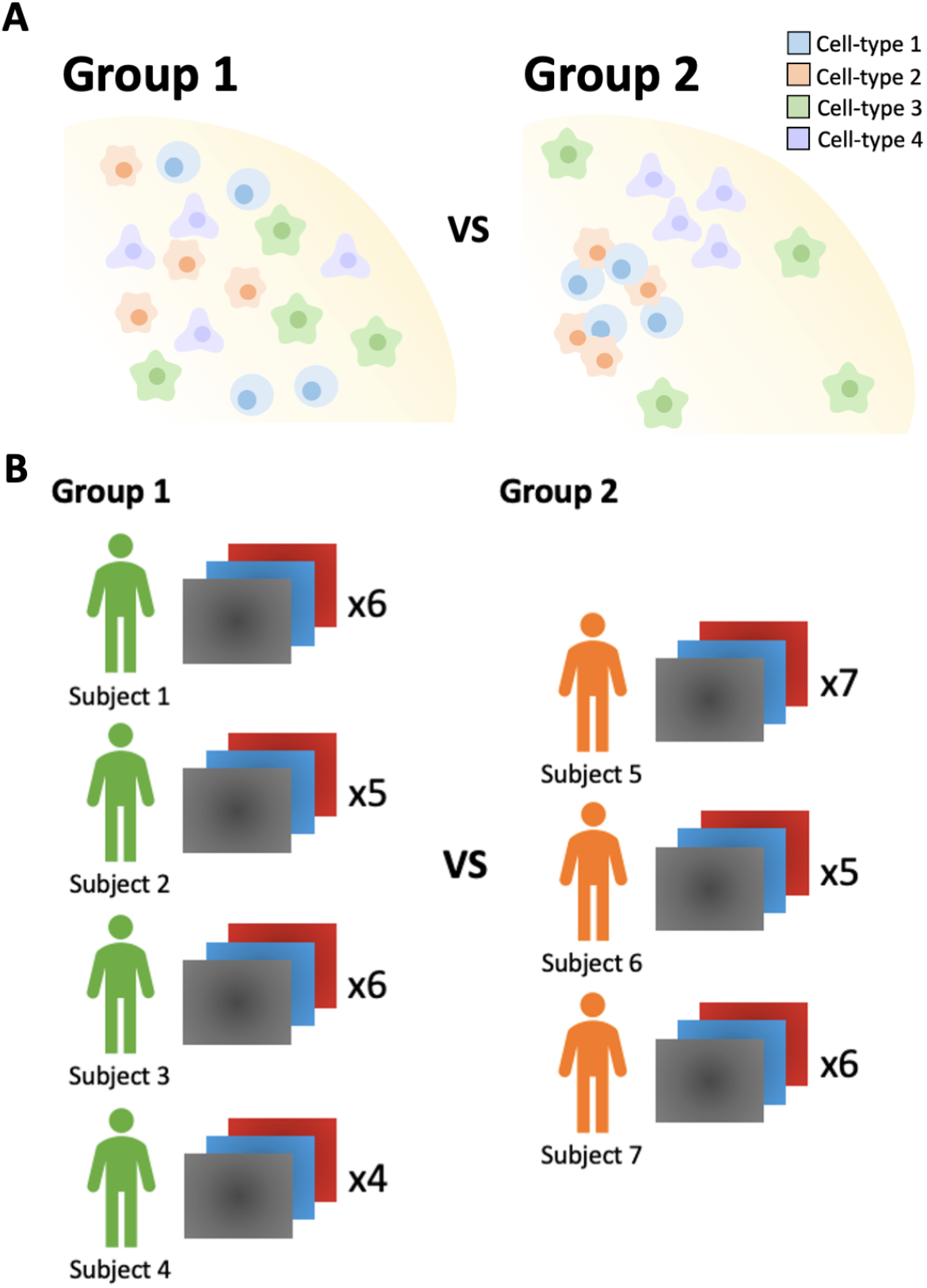
Schematic of the experimental motivation of spicyR. (A) Analytical question. From the images obtained, spicyR aims to identify differences in cell type localisations between the two groups. (B) Example experimental setup. Here, we have 4 subjects from group 1, and 3 donors from group 2, each with a different number of high parameter images. When applying spicyR, the number of subjects per group does not have to be equal. Likewise, the number of images per subject does not have to be equal.

### Construction of the L curve and localisation score

Following single-cell segmentation and classification, images are modelled as a ‘marked point process model’, in which each cell is represented as a point in a two-dimensional plane. Spatial localization between two cell types within an image can be quantified with a K-function (Ripley, 1976),

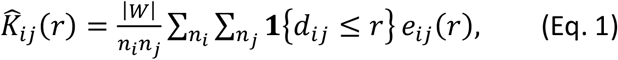

where 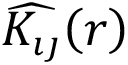 summarized the degree of localisation of cell type *j* with cell type *i*, *n*_*i*_ and *n*_*j*_ are the number of cells of type *i* and *j*, |*W*| is the image area, *d*_*ij*_ is the distance between two cells, and *e*_*ij*_(*r*) is an edge correcting factor. The K-function can be interpreted as the average number of cells of type *j* within a distance *r* away from each cell of type *i*.

The L-function is a variance stabilised K-function given by the equation

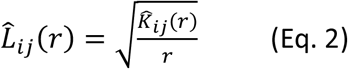

(Ripley, 1976). **Fig. 2B** shows an example observed L-curve compared to the expected L curve obtained from points distributed randomly in a Poisson distribution. Curves above the Poisson line can be interpreted as showing greater attraction when compared to a random distribution, and curves below the Poisson line can be interpreted as avoidance of the two cell types.

**Figure 2.**
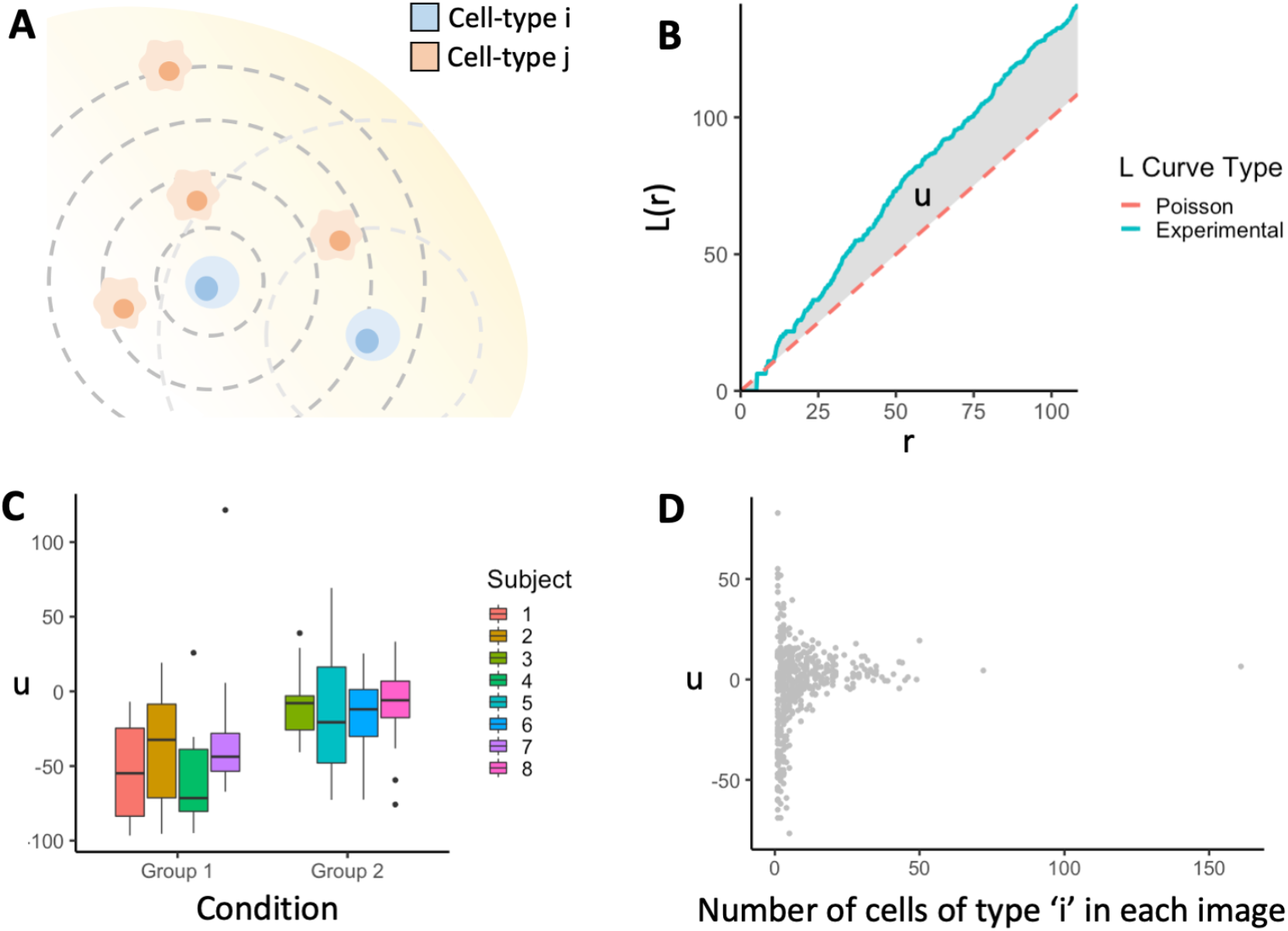
Summary of the spicyR framework. (A) Representation of two cell types in a 2D plane, which can be represented as points. Localisation of cell type j around cell type i can be modelled by using Ripley’s K-function. (B) Example of the experimental L curve (blue) compared against a Poisson L curve (red) expected if cell-type i and j were distributed independent of each other. The shaded area represents the AUC statistic used to assess whether localisation (positive value) or avoidance (negative value) is occurring. (C) A boxplot representation of the AUC statistics used to compare pairwise localisation across different groups. (D) Plot of the AUC statistic vs the number of cells of type i. As cell count is decreased, the variance of the AUC statistic increases. A generalized additive model is used to model the square of the statistic as a two-dimensional function of the counts of cell types ‘I’ and ‘j’, and these are used as weights in the linear models used.

To reduce these summary functions into a single localization score *u*, we take an area between curve measurement (**Fig. 2B**), given by the equation

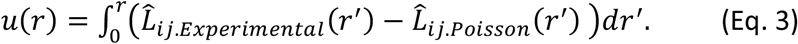

Here, values above zero represent greater attraction between cells when compared to random, while values below zero represent avoidance when compared to random.

### Weighting scheme used to account for varying cell counts

The relationship between cell count and variability of u is modelled with a generalized additive model (GAM) fitted with mgcv (Wood, 2001). We fit a GAM to the square of *u* with the counts of both cell types *i* and *j* as explanatory variables. The inverse of this fitted curve is used as a weighting scheme for each measurement in the model described in the following section – that is, lower weights are applied to images with lower cell counts.

### Hypothesis testing with a linear model

To assess changes in cell type localisation between groups, we implement either a linear model, or a linear mixed-effects model with random intercepts using the lme4 package (Bates et al., 2015). Here, *u*_*ij*_ is the localization score for subject *i* and image *j*, the treatment group or condition (*x*_*i*_) is modelled as a fixed effect with coefficient β and if a subject has multiple images the intercept (*α*_*i*_) is modelled as a random effect. Other covariates are included as W with coefficients *Г*.

#### Linear model

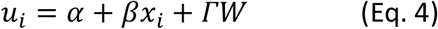

#### Linear mixed-effects model

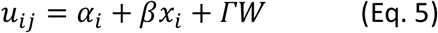

For the mixed effects model, p-values are calculated with Satterthwaite’s approximations using the lmertest package (Kuznetsova et al., 2017). Alternatively, we can also calculate p-values using bootstrapping in which the data is resampled and the model is reapplied. The p-value is calculated by comparing the distribution of fixed effects constants to zero; i.e.

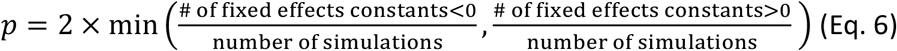

### Simulations to assess the performance of spicyR

To examine the performance of spicyR, simulations were generated using the spatstat package, as summarised in **Fig S2**. In summary, 10000 ‘null’ simulations were performed, which showed no cell localisation changes. 10000 simulations where there were differences in cell localisation were then performed.

Each simulation contained 20 subjects, each with two groups of 10 images. In both sets of simulations, the images in group 1 are generated such that it contains two cell types which do not show localisation. In the null simulations, group 2 is generated in the same way, while in the simulation with changes, group 2 is generated such that the two cell types are localising with each other. Hence, the null simulations simulate no cell localisation changes, while the simulations with changes simulate increased cell localisation in group 2 compared to group 1.

Images of size 1000 × 1000 units^2^ were generated. All group 1 images were generated using a Poisson point process model to simulate randomness. In the null simulations, group 2 images were also generated using a Poisson point process model. In the group 2 images of the simulations with changes, cell type A was generated with a Poisson point process model, while cell type B was distributed with a Poisson point process with a density weighting obtained from the distribution of cell type A. This simulated a distribution of cell type B that was dependent on the distribution of cell type A, thus simulating localisation. For each subject, variation was introduced by varying the density smoothing kernel size in the simulations with changes. Additionally, random cell counts were introduced for each image.

In summary, for each simulation 20 subjects were generated, each with two groups of 10 images. spicyR was then applied to each simulation to obtain a p-value from which the significance of the changes in cell localisation can be identified. ROC curves, as well as barplots of the percentage of simulations that were significant were obtained for both null and localisation simulations.

### Application to diabetes imaging mass cytometry data in Damond et al. 2019

As an implemented example of the framework, we apply spicyR to the data presented by Damond et al. (2019). This study aimed to identify the spatial distribution of markers and cells in pancreatic islets, comparing three different stages of type-1 diabetes mellitus (T1DM) progression: non-diabetic, onset diabetes, and long-duration diabetes, with four subjects per group. The single cell image data was downloaded from Mendeley (Version 2: https://data.mendeley.com/datasets/cydmwsfztj/2). This dataset consists of images of pancreatic islet cells across the three disease stages, obtained via imaging mass cytometry with 37 markers. The number of images per subject ranged from 64-81 (Total = 845; Non-diabetic = 274; Onset = 290; Long-duration = 281). Additionally, the dataset contained spreadsheets providing relevant patient information, as well as the cell coordinates, marker expression, and cell classification.

Here, we aimed to apply spicyR to identify significant differential localisations across the different disease stages. The X and Y co-ordinates of each cell, which image the cell belonged to, and the cell type were obtained, alongside the patient ID and diabetes disease stage for each image. Images belonging to the non-diabetic and onset diabetes groups were included, while the long-duration diabetes group was excluded. SpicyR was then applied to the data, with patient ID being treated as the random effect and the disease stage (non-diabetic vs onset diabetes) being treated as the fixed effect. The localisation statistic was obtained by integrating between a distance range of 0 to the default maximum value of spatstat (1/4 the size of the smallest edge of the image). Finally, the number of bootstrapping simulations was set to 20000 (minimum p-value of 0.0001). The cell types studied here are the endocrine cell subsets (alpha, beta, gamma, and delta) and the immune cell subsets (Naïve T, T helper, T cytotoxic, neutrophils, and macrophages).

## Results

Here we present spicyR, a framework for identifying changes in spatial association between pairs of cell types that could be associated with images from different clinical or experimental groups (**Fig. 1**). As input, spicyR requires images that have undergone single-cell segmentation and cell type classification. We provide functionality for empirically estimating the variability of the spatial associations between cell types within an image, for including multiple images per subject in the models and for both parametric and non-parametric estimates of p-values.

We observed across multiple datasets (**Fig. 2D**, **Sup Fig. 1**) that the variability of the measure of spatial association between a pair of cell types, *u,* decrease as the number of the cells in an image increase. Quantifying the relationship between the variability of *u* and the number of cells in an image, provides an opportunity to propagate this information in the model fitting process. We model the relationship between *u* and cell count by fitting a generalized additive model (using the mgcv package; (Wood, 2001)) to the square of *u* as a function of the counts of cell types (**Fig. 2D**). The inverse of the values obtained from this fitted curve are then used as weights when testing for association between the localisation of a pair of cells and an outcome, with lower weights being applied to images with lower cell counts.

Simulated images were generated using the spatstat package to assess the performance of spicyR (**Fig. S2)**. The generated ROC curve (**Fig. 3A**) showed that the weighted tests performed better than the non-weighted tests, with bootstrapping further improving performance slightly (AUC values: No Weights = 0.893; No Weights + Bootstrap = 0.901; Weights = 0.945; Weights + Bootstrap = 0.952). When a p-value cutoff of 0.05 was introduced, the percentage of true-positive interactions was highest in the bootstrapped weighted tests (**Fig. 3B**) and the false positive rate was closest to 5%. Hence, choosing to use weights that are inversely related to the variability of the localization score *u* with a bootstrap for calculating p-values will provide the best performance.

**Figure 3.**
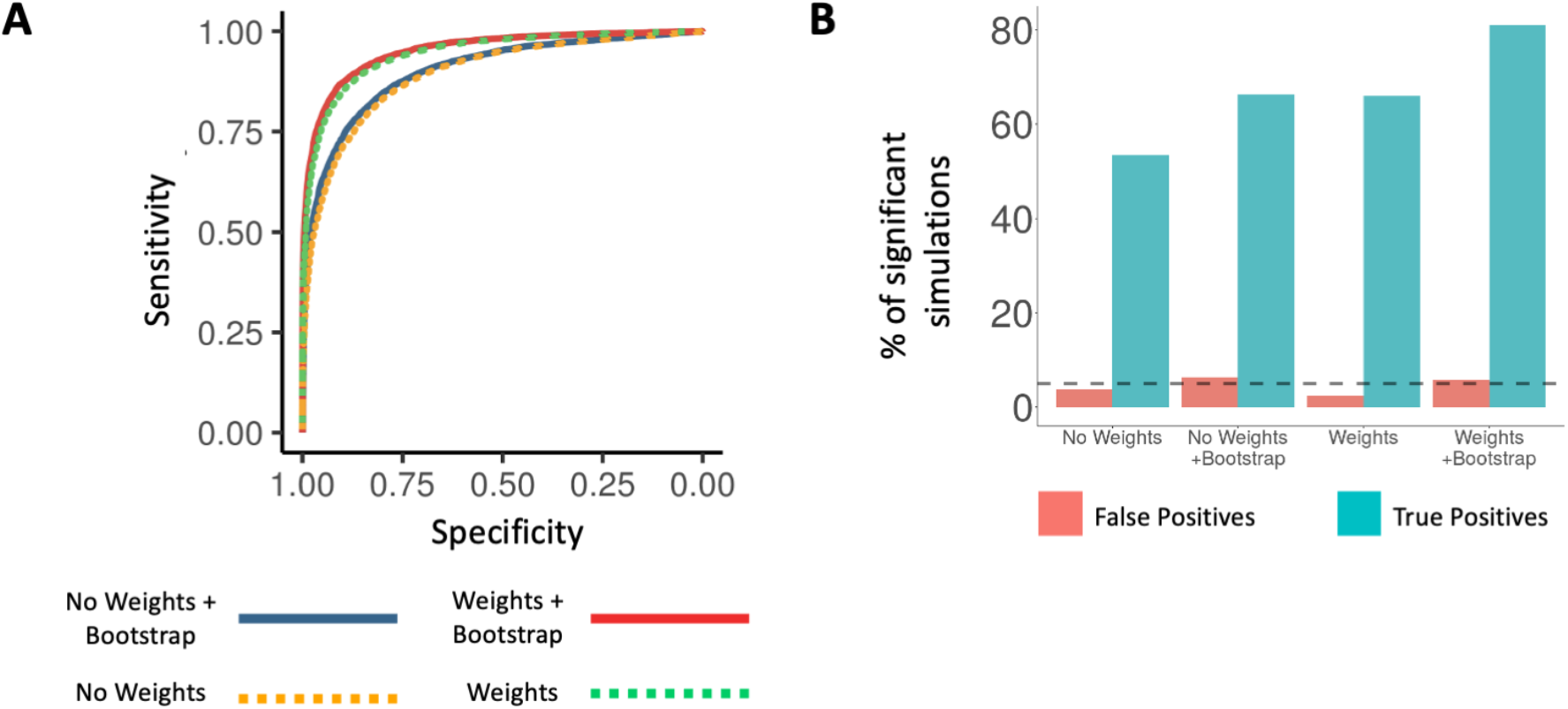
Simulations demonstrating the performance of the spicyR framework. Two sets of 1000 simulations were performed: the null simulations, generated without cell localisation differences between two groups, and simulations generated with changes in cell localisation between two groups. (A) ROC curves obtained from simulations (AUC: No Weights = 0.893; No Weights + Bootstrap = 0.901, Weights = 0.945; Weights + Bootstrap = 0.952). (B) Bar plots of the percentage of null simulations (false positives) and simulations with changes in localisation (true positives) that were significant at a p=0.05 level when spicyR was applied.

Damond et al. (2019) aimed to identify spatial changes in marker and cell distribution in pancreatic islets of three T1DM disease groups: non-diabetic, onset diabetes, and long-duration diabetes. A key finding of the study was that there was a temporal correlation between beta cell destruction, marked by marker loss and cell decreases, and an increased infiltration of T helper (Th) and T cytotoxic (Tc) cells in beta cell-rich pancreatic islets. Hence, Th and Tc cells were implicated in the destruction of beta cells characteristic of T1DM. We aimed to recapitulate this result, by showing that Th and Tc cells are more localised to beta cells in the onset diabetes group compared to the non-diabetic group.

We applied spicyR to identify differential cell type localisations between the non-diabetic group and the onset diabetes group. The cell types included here are the endocrine cell subsets (alpha, beta, gamma, and delta), and the immune cell subsets (Naïve T, T helper, T cytotoxic, neutrophils, and macrophages). **Fig. 4A** shows a heatmap representing the −log10 of the p-values obtained. It was found that Th and Tc cells showed increased spatial localisation with islet cells in the onset diabetes group, specifically towards beta cells. The T cell localisation was more significant with beta cells (p = 0.0001 for both Tc and Th cells) compared to alpha cells (Th to alpha: p = 0.0203; Tc to alpha: p = 0.1424). This suggests a migration of Th and Tc cells towards beta cells specifically during the early disease stages, reiterating the results presented by Damond et al. **Fig. 4B** shows representative masks reflecting these results. It was also found that there was increased localisation between Th cells, and between Th (p = 0.0001) and Tc (p = 0.0117) cells in the onset diabetes group. However, Tc cells did not show increased interactions with each other (p = 0.6622). Furthermore, there was an overall increase in immune-immune interactions: for example, there was an increased interaction between Th cells with B cells (p = 0.001), and between Tc cells and neutrophils (p.= 0.001). Hence, spicyR is able to find key changes in immune cell localisations with diabetes progression which was not highlighted by the original analysis. Finally, there was significant loss of localisation between beta cells with each other in the onset diabetes group. Again, this change in spatial architecture was not highlighted in the original publication. Overall, spicyR is able to identify significant changes in the spatial distribution of cells with diabetes progression.

**Figure 4.**
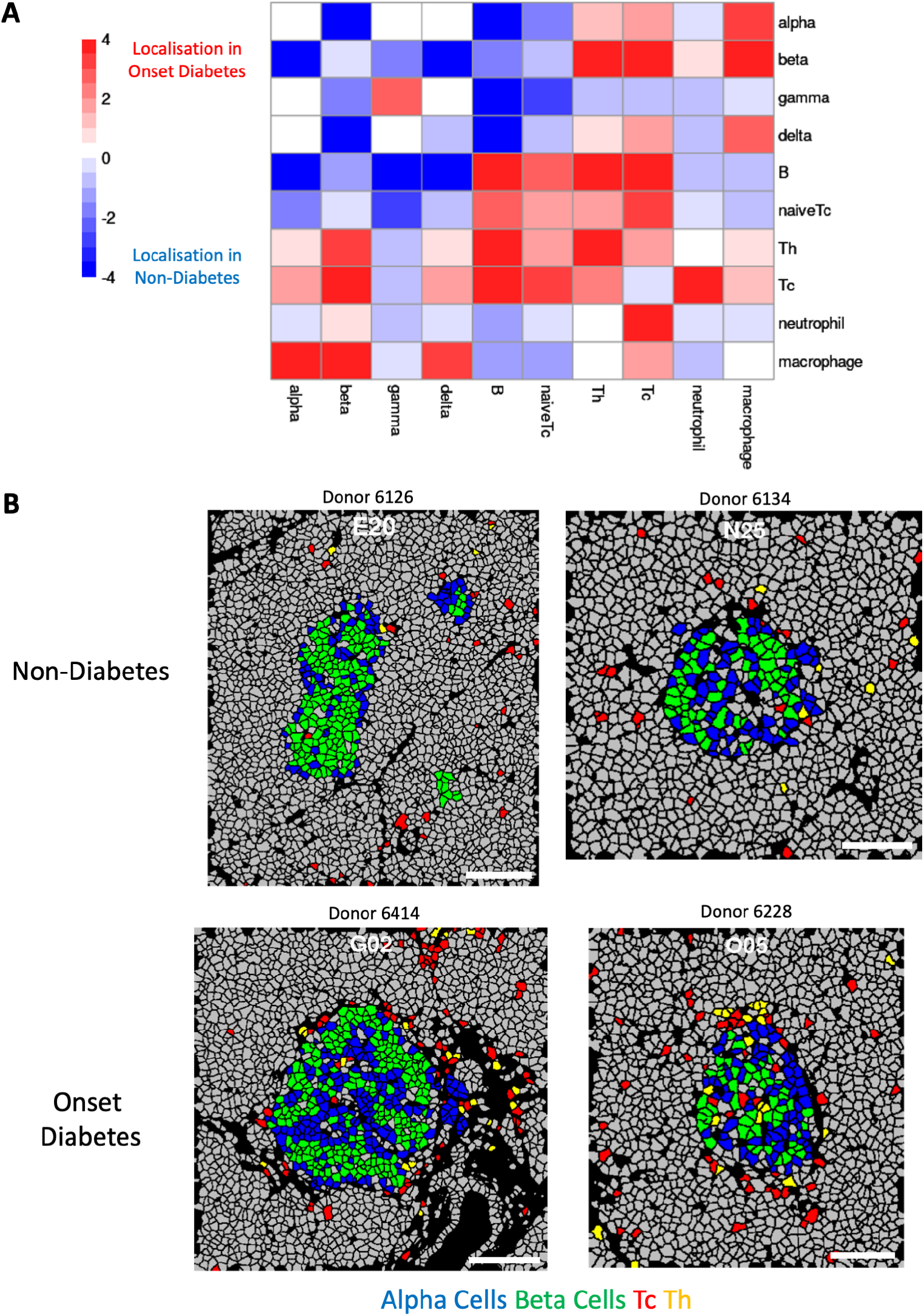
Application of spicyR to the Damond et al. (2019) type-1 diabetes IMC dataset. The spicyR framework was used to compare changes in cell localisations between the non-diabetic group and the onset diabetes group. (A) Heatmap showing the –log10 p-value when applying spicyR with the number of bootstrap simulations set to n = 20000. The y-axis represents cell type *i* (the ‘from’ cell type), and the x-axis represents cell type *j* (the ‘to’ cell type). Positive values (red) represent increased localisation in the onset diabetes group; negative values (blue) represent increased localisation in the non-diabetic group. (B) Masks showing representative images from the non-diabetes group and the onset diabetes group. Four cell types are highlighted in blue (Alpha cells), green (Beta cells), red (Tc) and yellow (Th).

## Discussion

Here we have presented spicyR, a tool for identifying differential cell localisations across different groups. We have demonstrated its performance to identify cell localisation changes through both simulated images, and with application to a type-1 diabetes IMC dataset. Simulations revealed that including weights that quantified the relationship between the number of cells in an image and the variability of a localisation statistic increased the sensitivity and specificity of spicyR. Furthermore, when spicyR was applied to the diabetes data set, the original results were reaffirmed and other key cell interactions present in diabetes progression were highlighted.

The spicyR package has many advantages compared to other differential localisation strategies used in high parameter image analysis. The key advantage of the package is its ability to summarise changes in cell localisations across groups of images. If multiple images are obtained from multiple subjects, the mixed effects model implemented allows variations within each subject to be modelled. By using an integrated spatial L curve, we obtain a localisation statistic that is easily interpretable and comparable across images. Furthermore, we implement a weighting scheme to account for variation in the localisation statistic given the pairwise cell type counts, which improves the predictive capabilities of spicyR. Finally, the package allows differential cell localisations to be identified across all pairwise cell types within the dataset, summarised in an interpretable heatmap. In this way, spicyR provides the framework for highlighting key cell-cell interactions that change across groups within a high parameter imaging dataset.

Despite the advantages discussed, there are several limitations to the implementation and evaluation of spicyR. Firstly, to benefit from the weighting regime, a moderate number of images are required to better model the relationship between cell count and localisation score. Depending on the experimental approach, and the availability of biological samples, this may not be feasible. Secondly, there are trade-offs to using a small vs a large radius when applying spicyR. A large radius may provide a better overall summary of cell-type localisation within an image, but can have decreased sensitivity, particularly if the localisation occurs strongly only over short distances. Hence, users should explore and choose an appropriate distance based on the biological questions being studied. Finally, while computational simulations were performed, it is difficult to validate spicyR within biological scenarios. There may be further scope for producing biologically relevant ground truths with which spicyR can be tested against, as well as more complicated simulation studies to elicit the effectiveness of the package.

It is important to acknowledge that spicyR should not be used in isolation to other analysis techniques, with the package being a key step in a broader analysis pipeline for high parameter imaging data. Firstly, spicyR is contingent on single-cell segmentation and classification being sufficiently accurate to elicit biologically relevant results. It is also complementary to other tests such as testing for changes in cell type composition and cell marker expression (Baharlou et al. 2019) which are necessary for providing an overview of the biological process being studied. Furthermore, it is advised to explore images visually both before and after any spatial analysis to identify whether visual observations appear to be consistent with the results of spicyR. Tools such as histoCAT (Schapiro et al. 2017) and the Bioconductor package cytomapper (Eling et al. 2020) provide useful exploratory tools for facilitating such exploratory data analysis. Ultimately, the package serves the role of highlighting changes in cell type localisation. This information can be crucial for synthesising key biological insight from multiplexed imaging experiments. Overall, results from spicyR will complement observations obtained from other elements of an image analysis pipeline.

## Acknowledgements

This work has been partly supported by the University of Sydney and an Australian Research Council Discovery Early Career Researcher Award (DE200100944) funded by the Australian Government. The funders had no role in the study design, data collection and analysis, decision to publish, or preparation of the manuscript.

---

**Figure S1.**
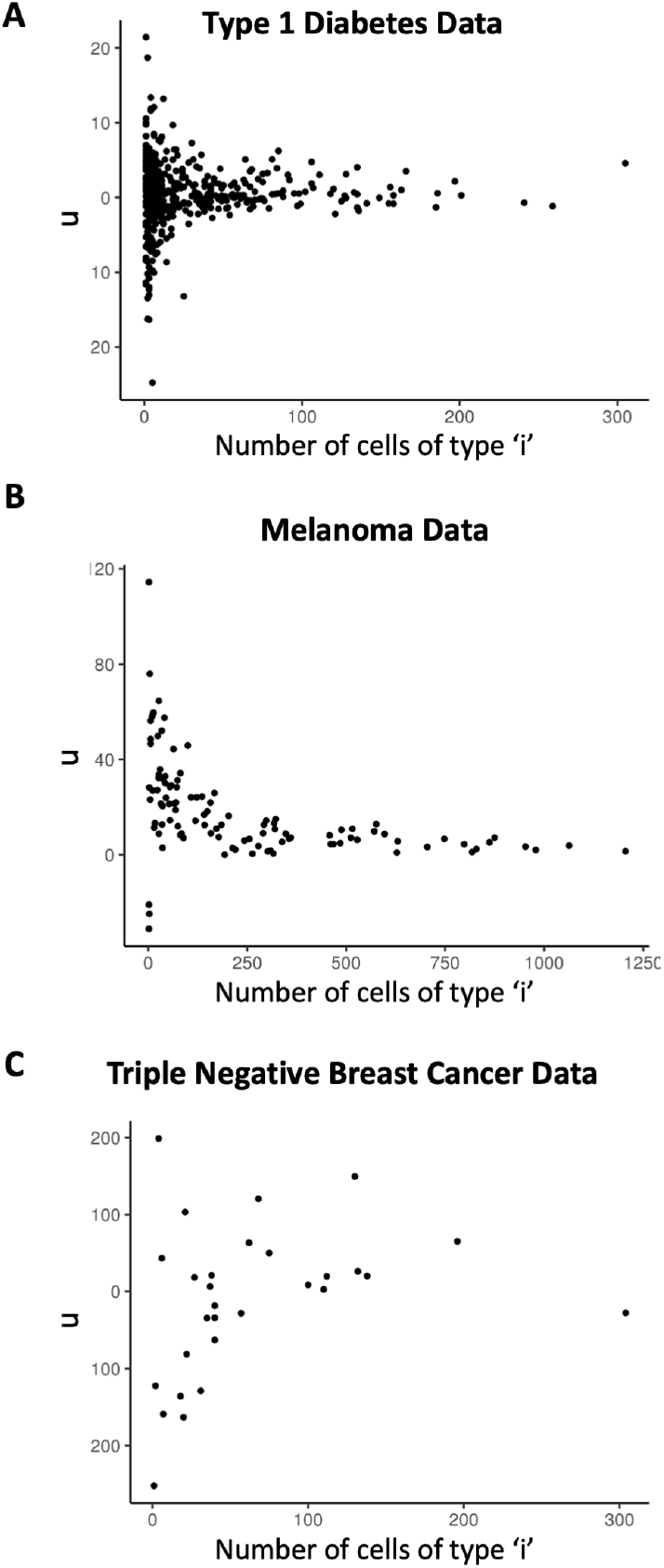
Relationship between the number of cells and variability of the quantification of localisation. u as a function of the number of cells of type ‘i’ for each image. As the number of cell types is decreased, the number variation in the u statistic is increased. This is seen across three different datasets: (A) Imaging mass cytometry images of type 1 diabetes samples from Damond et al. 2019 (B) Opal Multiplex immunohistochemistry images of melanoma samples from Willmott Lab (unpublished) (C) Multiplexed ion beam imaging by time-of-flight (MIBI-TOF) images of triple negative breast cancer samples from Keren et al. 2018.

**Figure S2.**
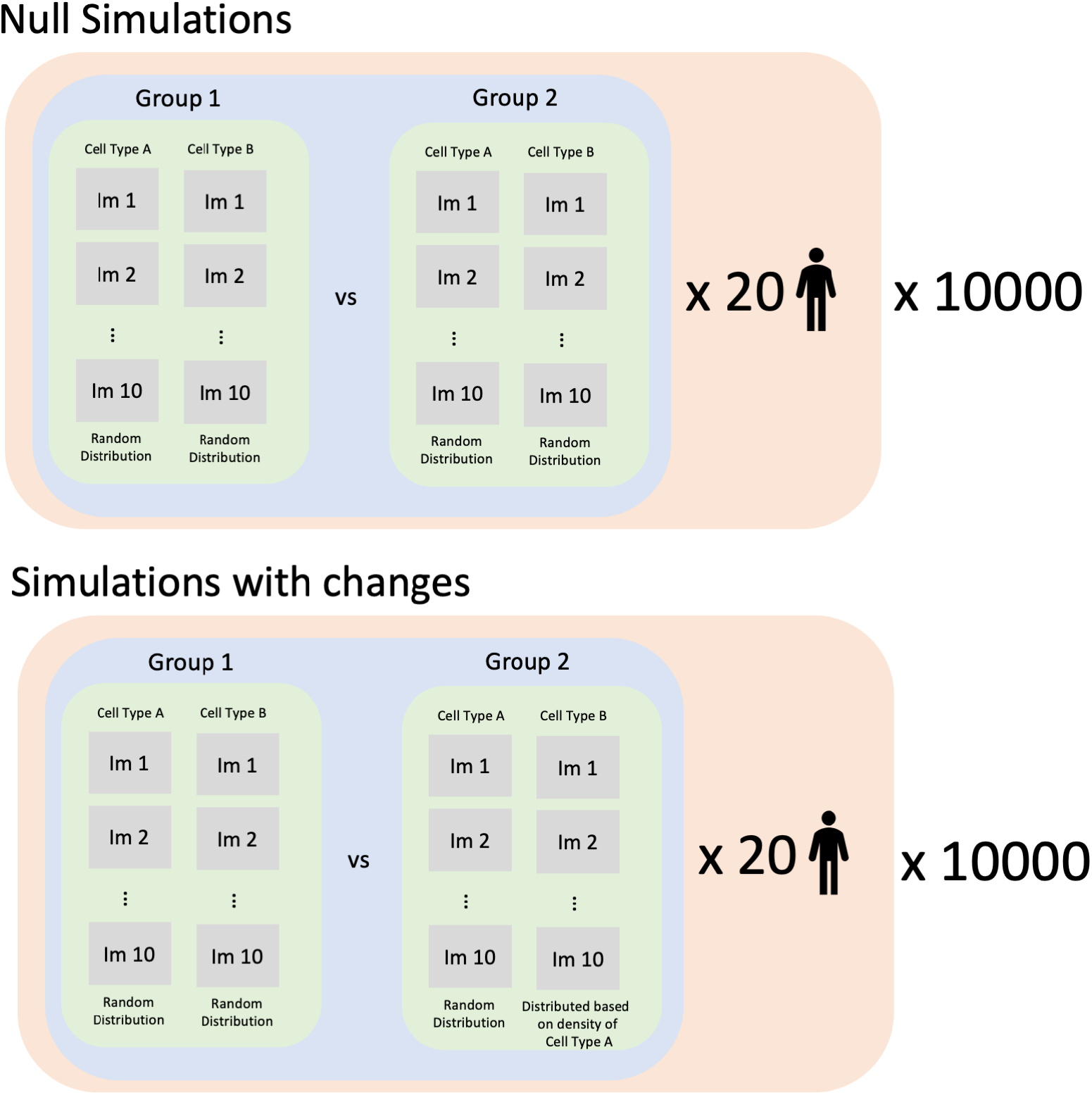
Summary of the simulations performed to assess the robustness of spicyR. 10000 null simulations and 10000 simulations with differences in cell localisations were performed. Each simulation contained 20 subjects, each with two groups of 10 images. Simulations were performed in such a way that the null simulations simulate no cell localisation changes, and the simulations with changes simulate increased cell localisation.

