## Supplementary Figures for "spicyR: Spatial analysis of *in situ* cytometry data in R"

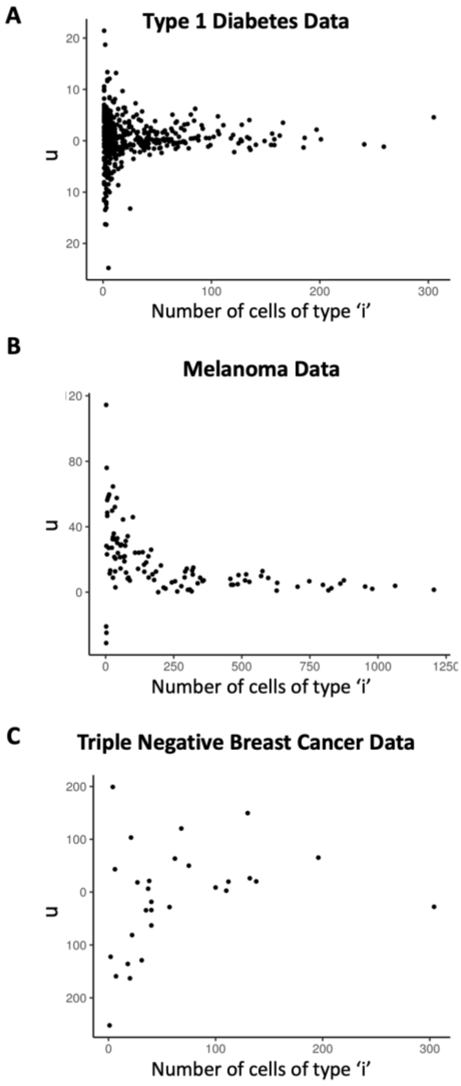


**Figure S1. Relationship between the number of cells and variability of the quantification of localisation.**

u as a function of the number of cells of type ‘i’ for each image. As the number of cell types is decreased, the number variation in the u statistic is increased. This is seen across three different datasets:

(A) Imaging mass cytometry images of type 1 diabetes samples from Damond et al. 2019

(B) Opal Multiplex immunohistochemistry images of melanoma samples from Willmott Lab (unpublished)

(C) Multiplexed ion beam imaging by time-of-flight (MIBI-TOF) images of triple negative breast cancer samples from Keren et al. 2018.

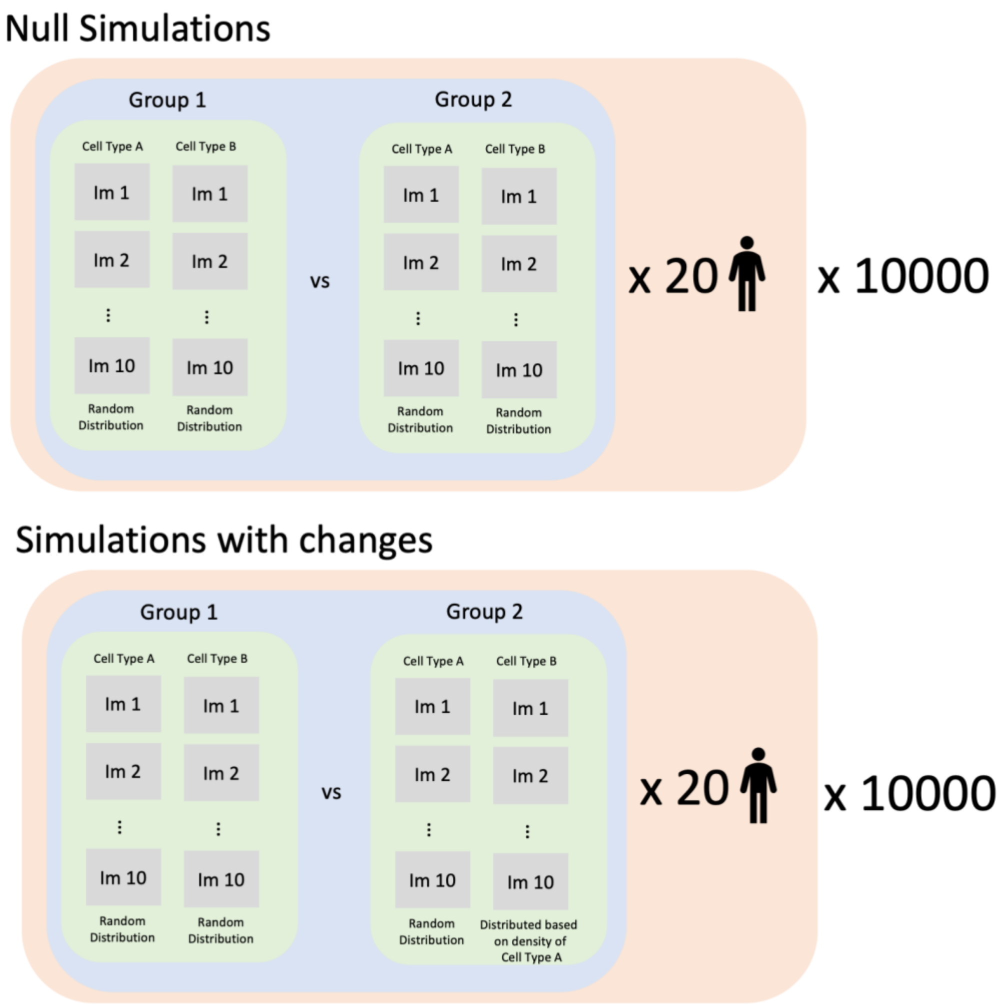


**Figure S2. Summary of the simulations performed to assess the robustness of spicyR**

10000 null simulations and 10000 simulations with differences in cell localisations were performed. Each simulation contained 20 subjects, each with two groups of 10 images. Simulations were performed in such a way that the null simulations simulate no cell localisation changes, and the simulations with changes simulate
